# Two different adaptive speciation mechanisms operate during adaptation to a novel hot environment

**DOI:** 10.1101/2021.11.08.467720

**Authors:** Sheng-Kai Hsu, Wei-Yun Lai, Johannes Novak, Felix Lehner, Ana Marija Jakšić, Elisabetta Versace, Christian Schlötterer

## Abstract

**Background:** Ecological speciation and mutation-order speciation are two different mechanisms of adaptation-driven speciation. Both mechanisms predict different patterns of reproductive isolation for replicate populations adapting to the same environment. With ecological speciation, barriers to gene flow emerge between populations from different environments, but not among replicate populations from the same environment. Mutation-order speciation predicts reproductive isolation among populations adapted to the same environment.

**Results:** We demonstrate that both speciation processes occurred within about 100 generations when replicate *Drosophila simulans* populations adapted to a novel, hot environment. Gene expression analysis identified the underlying molecular mechanisms. Premating ecological speciation is the byproduct of an altered lipid metabolism, which also changed the cuticular hydrocarbon (CHC) composition in hot-evolved flies. Postmating reproductive isolation supports mutation-order speciation most likely driven by co-evolution of reproduction-associated genes.

**Conclusion:** Adaptation processes can rapidly induce incipient speciation and different speciation mechanisms affect pre- and postmating reproductive isolation. We propose that the definition of mutation-order speciation should be expanded to account for polygenic processes from standing genetic variation.

## Introduction

Most new species are formed by selection [1–3] as already suggested by Darwin [4]. Two different mechanisms by which selection can lead to speciation have been proposed, ecological speciation and mutation order speciation. During ecological speciation reproductive isolation occurs as a consequence of adaptation to different ecological stressors. Importantly, reproductive isolation is not the direct target of selection, but rather occurs because the selected genes either have pleiotropic effects on reproduction or are linked to another loci affecting reproduction [5]. Mutation-order speciation describes the random occurrence of different favorable, but incompatible, mutations in distinct populations adapting to the same environmental stressor [6, 7]. Many examples of ecological speciation are documented [e. g. 8, 9-12] and also mutation order speciation has received some empirical support [13].

Both speciation mechanisms make different predictions for populations evolving independently in the same environments. Ecological speciation predicts that reproductive isolation develops between populations adapted to different environments, but not among different populations in the same environment [2]. With mutation-order speciation, independent populations which are adapted to the same environment become incompatible because they use different sets of incompatible mutations. Mutation-order speciation can result from intra-genomic conflict as well as antagonistic co-evolution between the two sexes [2].

Experimental evolution provides an excellent empirical approach to study speciation processes, because environmental conditions can be well-controlled and replicate populations are exposed to the same environmental stress. Nevertheless, only ecological speciation has received strong empirical support by laboratory experiments [14–18] and in most cases the selected traits were not identified.

Here, we take advantage of a highly replicated experimental evolution study, where 10 replicate populations derived from a single polymorphic *D. simulans* population have adapted independently to a novel hot temperature regime in about 100 generations [19, 20]. Although ambient temperature is one of the major ecological factors driving adaptation in natural populations, it is not clear from previous work to what extent temperature has triggered speciation processes [21]. We find support for ecological speciation, because premating isolation occurred between ancestral and hot-evolved populations, but no difference was found among the replicates, which evolved independently in the same novel hot environment. Consistent with the predictions of mutation order speciation, we observed postmating reproductive isolation between the evolved replicates. We propose that ecological speciation is based on a modified lipid metabolism, which also causes shifts in CHC composition, a well-understood mechanism of reproductive isolation in fruit flies [22–24]. Because mutation-order speciation can involve inter-genic antagonistic co-evolution [2], we suggest that the high variance in evolutionary expression change of reproduction related genes drives the observed post-mating reproductive isolation.

## Results

After more than 100 generations of adaptation to a novel hot temperature regime, we estimated reproductive isolation (Figure 1). As predicted for ecological speciation, we observed significant mate discrimination between ancestral and hot-evolved populations but not for replicate populations evolved independently to the same selection regime. While hot-evolved males were more actively chasing females than ancestral males, the courting behavior was significantly reduced when exposed to ancestral females. The same pattern was seen for males from the ancestral population (Figure 2a, Kruskal-Wallis test, p < 0.05). Reproductive isolation was further shown in an assortative mating experiment where flies from two populations were mixed and mating partners could be chosen freely. Combining ancestral and evolved populations resulted in strong positive assortative mating, while no assortative mating could be detected among independently evolved populations (Figure 2b). These patterns of premating reproductive isolation are fully consistent with the expectations for ecological speciation: significant differences between flies adapted to different environments, but no reproductive isolation between flies from different populations evolved in the same environment.

**Figure 1.**
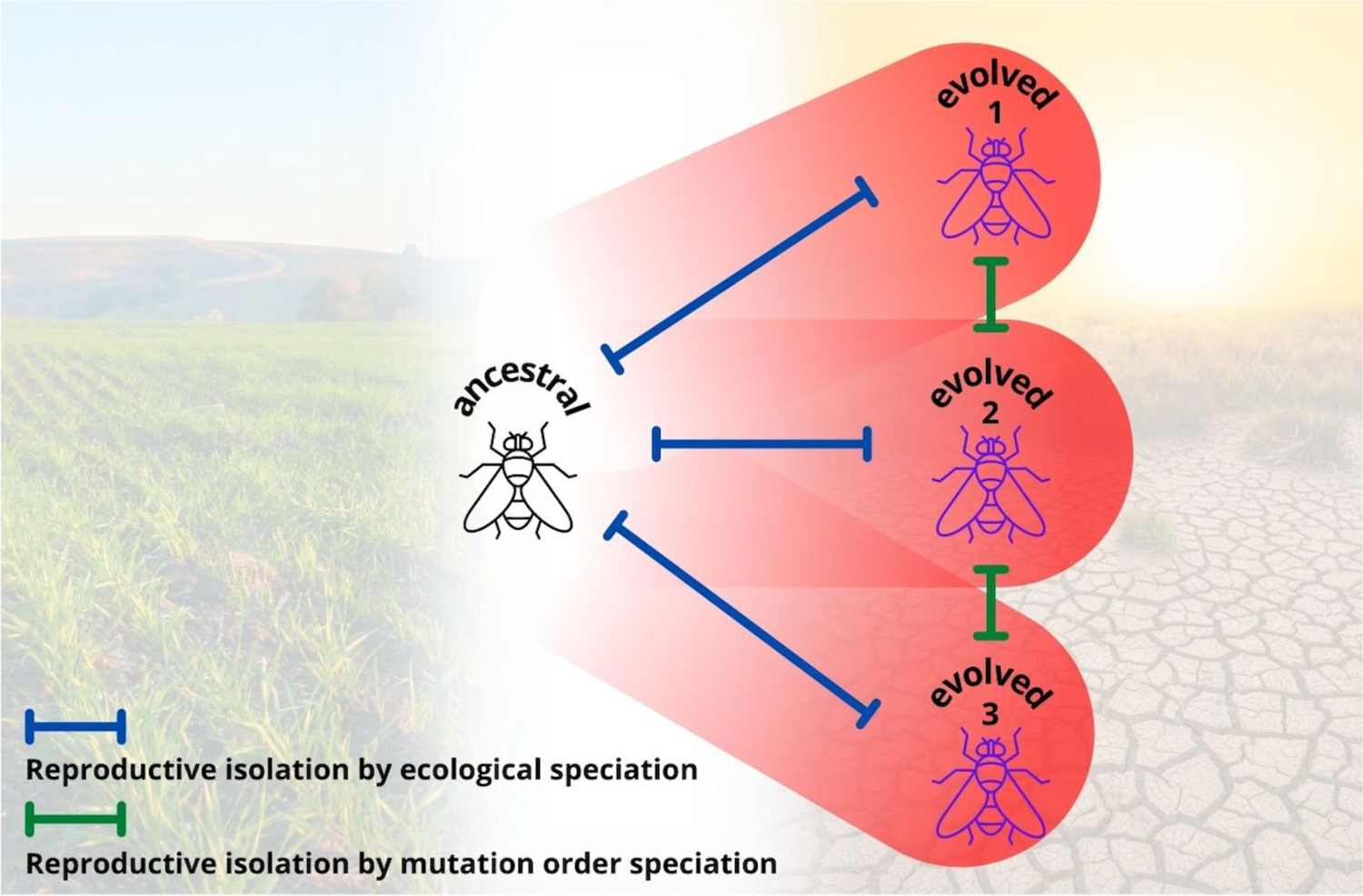
Schematic overview of the emergence of reproductive isolation during more than 100 generations of adaptation to a new hot environment. Replicate populations evolved at a hot temperature regime fluctuating between 18 and 28°C to mimic day and night conditions. Reproductive isolation evolved between ancestral and evolved population (ecological speciation) and among evolved replicates (mutation order speciation).

**Figure 2.**
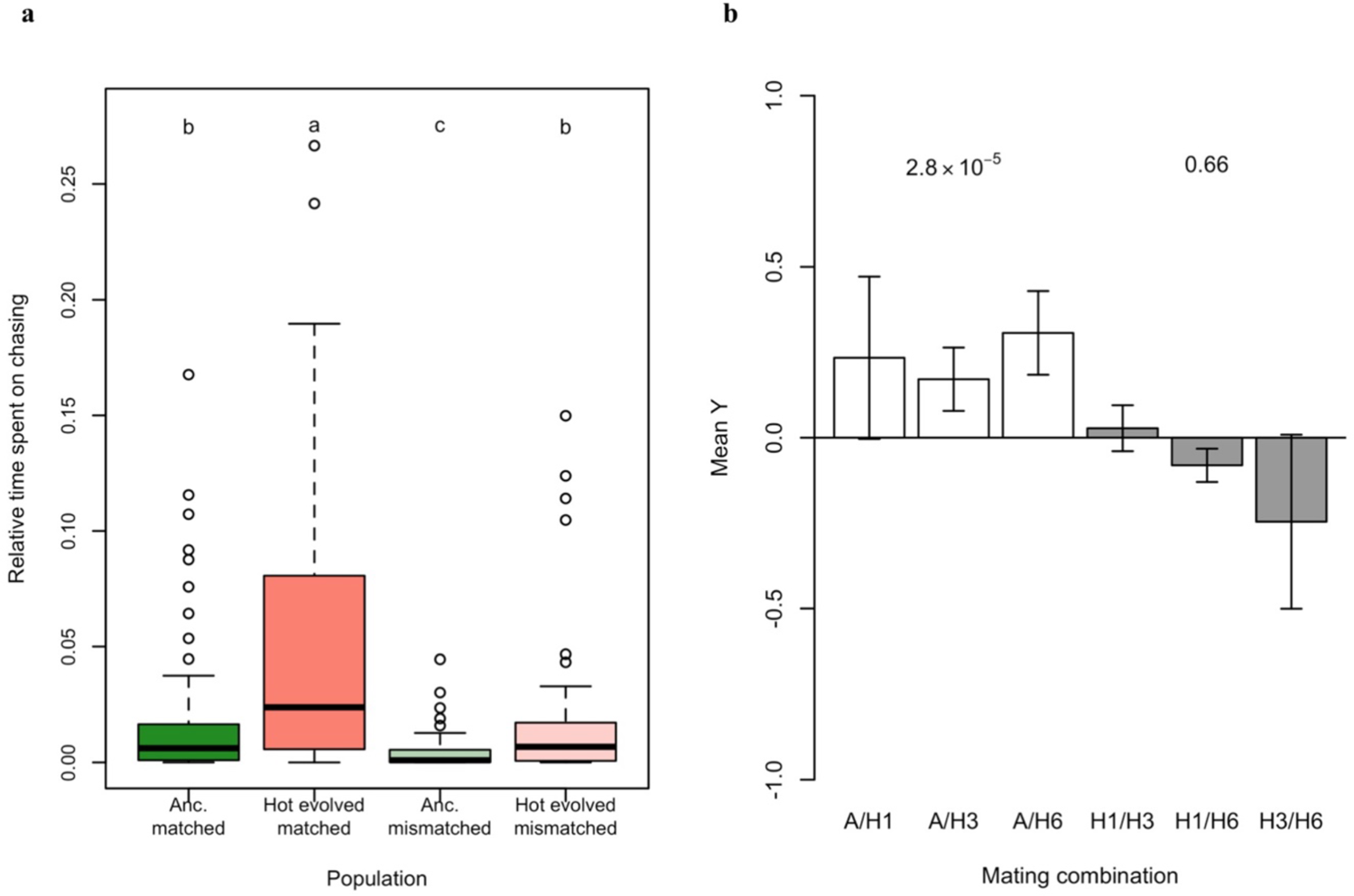
Evolution of mating preference and pre-mating reproductive isolation. **a.** Male reproductive activity in the presence of the females from the same evolved population or the ancestral one. The reproductive activity was measured as the time spent on chasing. Males spent significantly more time chasing females from the same populations (Kruskal-Wallis test, p < 0.001; post-hoc groups were indicated as the small-case letter above the boxes). **b.** Yule’s index for assortative mating. There is significantly positive assortative mating between flies from the same populations (white bars) while there’s no significant effect among the replicate evolved populations (p-values for Fisher’s exact test).

Previously we showed that the lipid metabolism is significantly altered in the hot evolved populations studied here [19]. Since the synthesis of cuticular hydrocarbons (CHCs), which serve a central role in mate recognition [22–24], is controlled by genes involved in lipid metabolism [25], we reasoned that the CHC composition may have been altered as a byproduct of the change in lipid metabolism. We tested the divergence in CHC composition between ancestral and evolved populations by measuring all hydrophobic compounds on the outer skeleton of virgin flies of each sex with gas chromatography/MASS spectrometry (GC/MS). We detected 18 major CHC compounds across both sexes, which differ in length of carbon chain or numbers of double bonds [22, 25] (Table 1 and Supplementary Figure 1). Principal component analysis (PCA) indicated that the relative abundance of the CHCs differed significantly between sexes as well as between evolved and ancestral populations (Figure 3a). The first PC, which explains 68.4% of total variance, reflects the differences between the two sexes – males synthesize more unsaturated CHCs than females (Figure 3b). The variation caused by adaptation to a hot laboratory environment is reflected by PC2 which accounts for 16.38% of the total variance (Figure 3a). No significant interactions between sexual dimorphism and evolution were observed for these traits (Table 1), indicating parallel evolution of CHC composition in both sexes. Evolved flies synthesize more unsaturated CHCs with longer carbon chains (Figure 3b and c). Interestingly, similar modifications in CHC compositions have been documented for multiple *Drosophila* species in latitudinal clines [26], implying that the same causal link with temperature adaptation is also present in natural populations.

**Figure 3.**
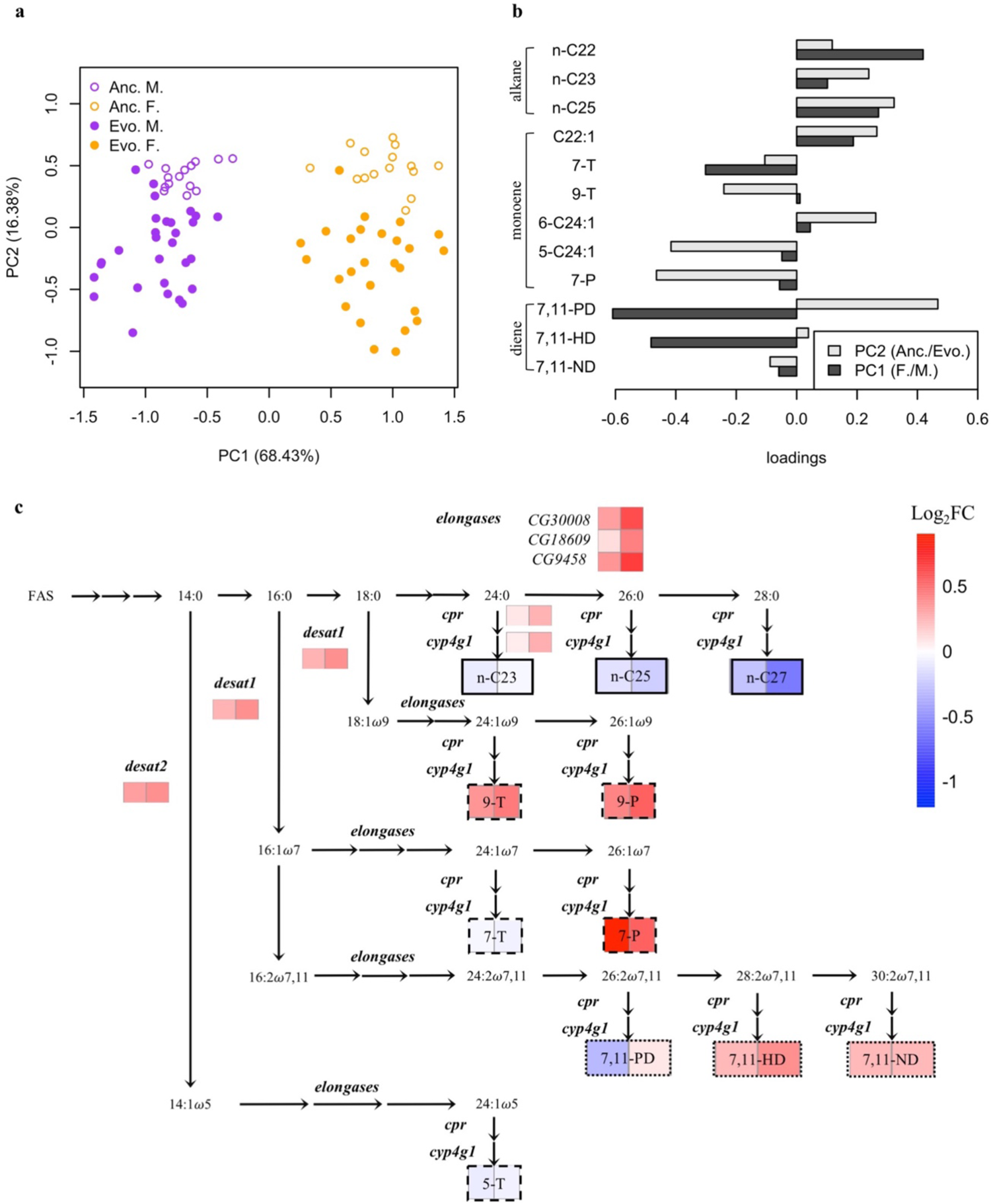
Evolution of cuticle hydrocarbons (CHCs) metabolism. **a.** Principal component analysis on the center-log transformed relative abundance of 12 CHCs detected in all samples. (orange: females; purple: males). Solid dots are evolved samples and empty ones are ancestral samples. **b.** Loading of each compound on the first and second principal components (PCs). Dark grey bars indicate the loadings for PC1. Positive loading suggests higher abundance in females. Light grey bars indicate the loadings for PC2 and positive values reflect higher abundance in the ancestral samples. **c.** Evolutionary changes in gene expression of the biosynthetic pathway for *Drosophila* cuticular hydrocarbons (CHCs). The heat maps underneath each compound (boxes with regular fonts) or genes (boxes with italic fonts) denote the changes in abundance during adaptation (log2-scale). The left cell denotes the changes in females and right cell for males. Red indicates increase and blue indicates decrease in abundance.

**Table 1.**
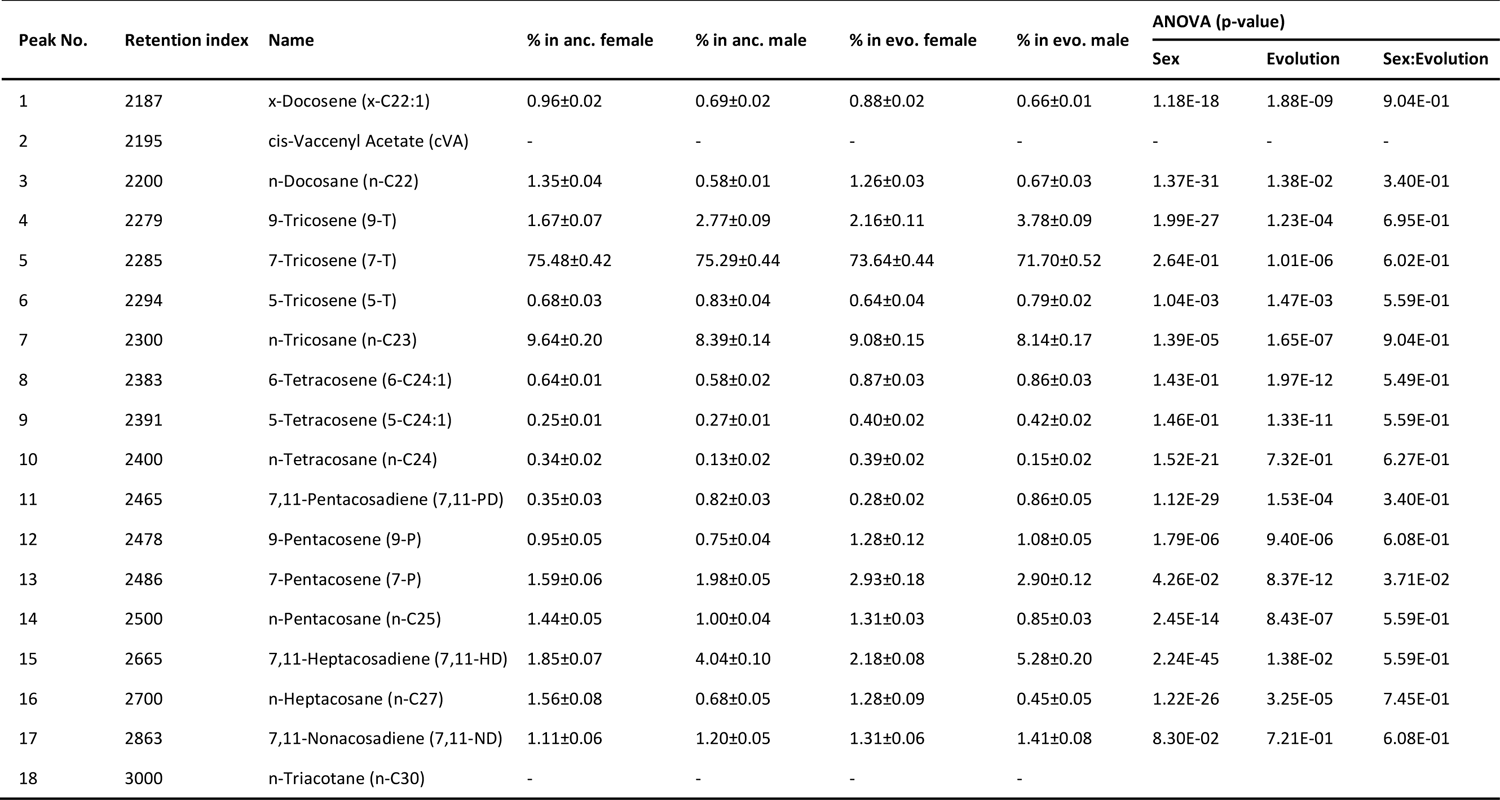
Abundance of cuticular hydrocarbons (CHCs) in relation to sex and evolution

| Peak No. | Retention index | Name | % in anc. female | % in anc. male | % in evo. female | % in evo. male | ANOVA (p-value) |  |  |
| --- | --- | --- | --- | --- | --- | --- | --- | --- | --- |
|  |  |  |  |  |  |  | Sex | Evolution | Sex:Evolution |
| 1 | 2187 | x-Docosene (x-C22:1) | 0.96±0.02 | 0.69±0.02 | 0.88±0.02 | 0.66±0.01 | 1.18E-18 | 1.88E-09 | 9.04E-01 |
| 2 | 2195 | cis-Vaccenyl Acetate (cVA) | - | - | - | - | - | - | - |
| 3 | 2200 | n-Docosane (n-C22) | 1.35±0.04 | 0.58±0.01 | 1.26±0.03 | 0.67±0.03 | 1.37E-31 | 1.38E-02 | 3.40E-01 |
| 4 | 2279 | 9-Tricosene (9-T) | 1.67±0.07 | 2.77±0.09 | 2.16±0.11 | 3.78±0.09 | 1.99E-27 | 1.23E-04 | 6.95E-01 |
| 5 | 2285 | 7-Tricosene (7-T) | 75.48±0.42 | 75.29±0.44 | 73.64±0.44 | 71.70±0.52 | 2.64E-01 | 1.01E-06 | 6.02E-01 |
| 6 | 2294 | 5-Tricosene (5-T) | 0.68±0.03 | 0.83±0.04 | 0.64±0.04 | 0.79±0.02 | 1.04E-03 | 1.47E-03 | 5.59E-01 |
| 7 | 2300 | n-Tricosane (n-C23) | 9.64±0.20 | 8.39±0.14 | 9.08±0.15 | 8.14±0.17 | 1.39E-05 | 1.65E-07 | 9.04E-01 |
| 8 | 2383 | 6-Tetracosene (6-C24:1) | 0.64±0.01 | 0.58±0.02 | 0.87±0.03 | 0.86±0.03 | 1.43E-01 | 1.97E-12 | 5.49E-01 |
| 9 | 2391 | 5-Tetracosene (5-C24:1) | 0.25±0.01 | 0.27±0.01 | 0.40±0.02 | 0.42±0.02 | 1.46E-01 | 1.33E-11 | 5.59E-01 |
| 10 | 2400 | n-Tetracosane (n-C24) | 0.34±0.02 | 0.13±0.02 | 0.39±0.02 | 0.15±0.02 | 1.52E-21 | 7.32E-01 | 6.27E-01 |
| 11 | 2465 | 7,11-Pentacosadiene (7,11-PD) | 0.35±0.03 | 0.82±0.03 | 0.28±0.02 | 0.86±0.05 | 1.12E-29 | 1.53E-04 | 3.40E-01 |
| 12 | 2478 | 9-Pentacosene (9-P) | 0.95±0.05 | 0.75±0.04 | 1.28±0.12 | 1.08±0.05 | 1.79E-06 | 9.40E-06 | 6.08E-01 |
| 13 | 2486 | 7-Pentacosene (7-P) | 1.59±0.06 | 1.98±0.05 | 2.93±0.18 | 2.90±0.12 | 4.26E-02 | 8.37E-12 | 3.71E-02 |
| 14 | 2500 | n-Pentacosane (n-C25) | 1.44±0.05 | 1.00±0.04 | 1.31±0.03 | 0.85±0.03 | 2.45E-14 | 8.43E-07 | 5.59E-01 |
| 15 | 2665 | 7,11-Heptacosadiene (7,11-HD) | 1.85±0.07 | 4.04±0.10 | 2.18±0.08 | 5.28±0.20 | 2.24E-45 | 1.38E-02 | 5.59E-01 |
| 16 | 2700 | n-Heptacosane (n-C27) | 1.56±0.08 | 0.68±0.05 | 1.28±0.09 | 0.45±0.05 | 1.22E-26 | 3.25E-05 | 7.45E-01 |
| 17 | 2863 | 7,11-Nonacosadiene (7,11-ND) | 1.11±0.06 | 1.20±0.05 | 1.31±0.06 | 1.41±0.08 | 8.30E-02 | 7.21E-01 | 6.08E-01 |
| 18 | 3000 | n-Triacotane (n-C30) | - | - | - | - |  |  |  |

We used gene expression data of the focal populations [19] to elucidate the basis of the CHC composition changes. Among the 469 genes with a significant increase in expression across replicate evolved populations, we found a prominent enrichment for genes involved in “very long-chain fatty acid biosynthetic process” in both sexes. 27 out of the 36 (33 are detected in our data) previously reported CHC synthesis-related genes in *Drosophila melanogaster* [25] showed significant expression changes in at least one sex during adaptation in the replicated *D. simulans* populations (Fisher’s exact test, odds ratio = 3.33, p-value < 0.01; Supplementary Figure 2). 16 of these genes changed their expression in the same direction in both sexes. *Desaturases* (*desat 1* and *desat 2*) and *elongases* (*CG9458*, *CG18609* and *CG30008*) were consistently up-regulated in all replicate evolved populations (Figure 3c and Supplementary Figure 2). This consistent transcriptional modification of long-chain fatty acid/CHC biosynthesis during the evolution can explain the significant changes in CHC composition in the evolved flies.

Reproductive isolation can occur before and after mating. While the significant premating isolation between ancestral and hot evolved populations supports ecological speciation, it is not informative about mutation-order speciation. Because most of the mechanisms proposed for mutation-order speciation operate after mating, we also tested postmating reproductive isolation among evolved replicates. F1 flies from crosses between replicates of the evolved populations produced 8.3% fewer viable offspring than F1 individuals from the same replicates of the evolved populations (Figure 4b, Wilcoxon’s test, p = 0.031). Considering the absence of pre-mating isolation (Figure 2b), the reduced hybrid fitness indicates that another (post-mating; fertilization- and/or viability-affecting) process occurs between the ancestral and evolved populations and leads to reproductive isolation.

**Figure 4.**
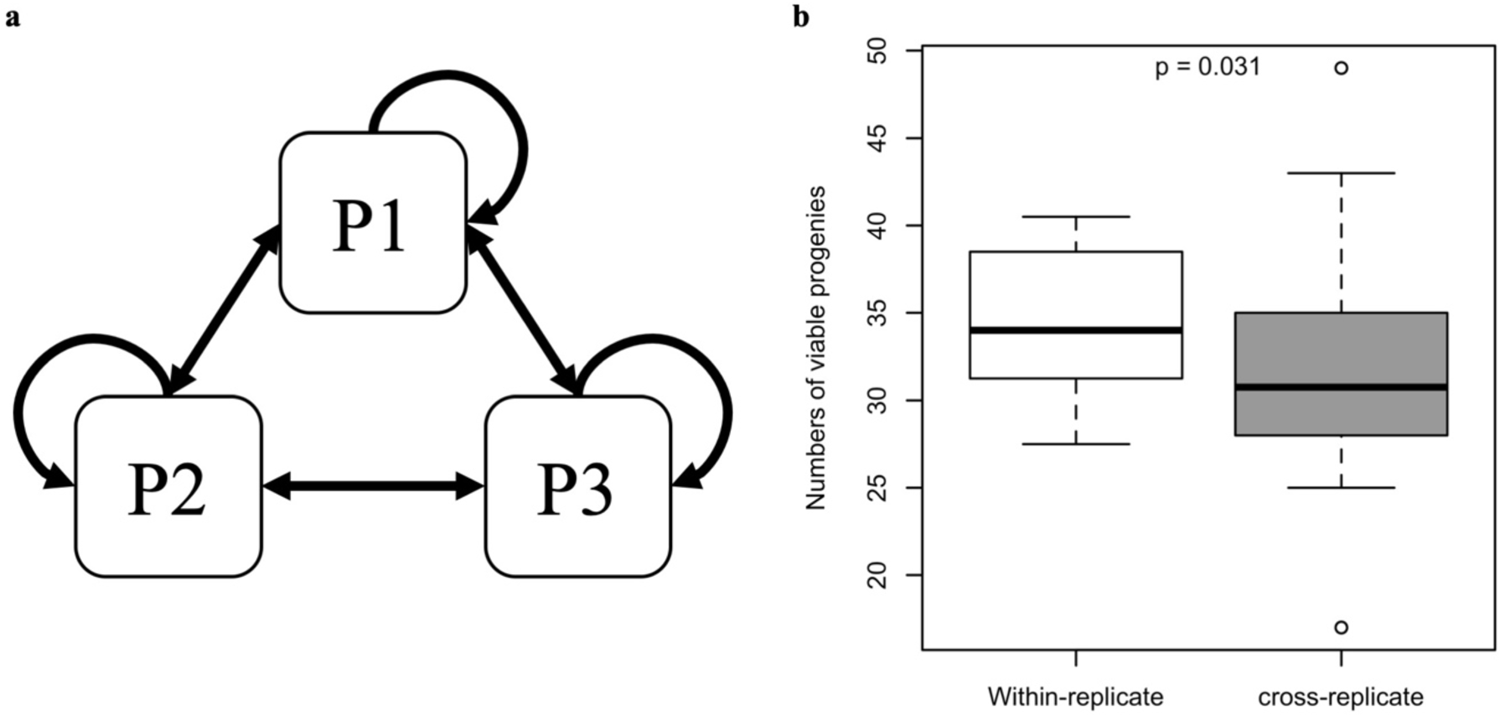
Evolution of post-mating incompatibility among independently evolved populations. **a.** Diallel cross design among three replicate evolved populations. Flies from each population were crossed with flies from the same population and the other populations reciprocally. **b.** Fitness difference between within-replicate crosses and cross-replicate ones. The fitness was measured as the numbers of viable adults in each cross. Fitness was significantly higher in within-replicate crosses (p = 0.031, Wilcoxon’s test), indicating post-mating incompatibility between independently evolved replicates.

We used RNA-Seq to explore the functional basis of this post-mating reproductive isolation. Reasoning that putative causal genes may diverge in gene expression, we searched for genes differentially expressed among replicate evolved populations. In total, we identified 3,062 genes which differed in at least one evolved population from the others (Figure 5 — Supplementary Figure 3, Supplementary file 1). Functional enrichment analysis revealed that a substantial portion of these divergently expressed genes is highly expressed in testis and is involved in reproduction-related processes (e.g. multicellular organism reproduction (GO:0032504); Supplementary file 2). Focusing on 255 reproduction-associated genes that diverged in expression across evolved replicated populations (see Materials and Methods), replicate-specific selection responses were detected (Figure 5a). The heterogeneity in gene expression evolution of reproduction-related genes was significantly higher than for a random set of genes (Figure 5b, Wilcoxon’s test, p < 0.001). Most of these reproduction-associated genes changed their expression in only one or two replicate population(s) towards the same direction (mostly up-regulation) during the adaptation (Figure 5a and Supplementary Figure 4, and Supplementary file 2). Hence, reproduction related genes are probably under selection in the new environment, but different sets of genes respond in different replicate populations. Given that mutation-order speciation can be mediated by inter-locus antagonistic coevolution [2, 27], our results provide a very compelling functional explanation to our observation: reproduction-associated genes are known to be expressed in males and females and their interaction is important for reproductive success [28, 29]. This implies that changes in gene expression in one replicate probably require adjustments of other genes to maintain functionality. The fitness decrease of hybrids between two independently evolved populations may be explained by a perturbation of the co-evolved gene expression patterns of reproduction-associated genes.

**Figure 5.**
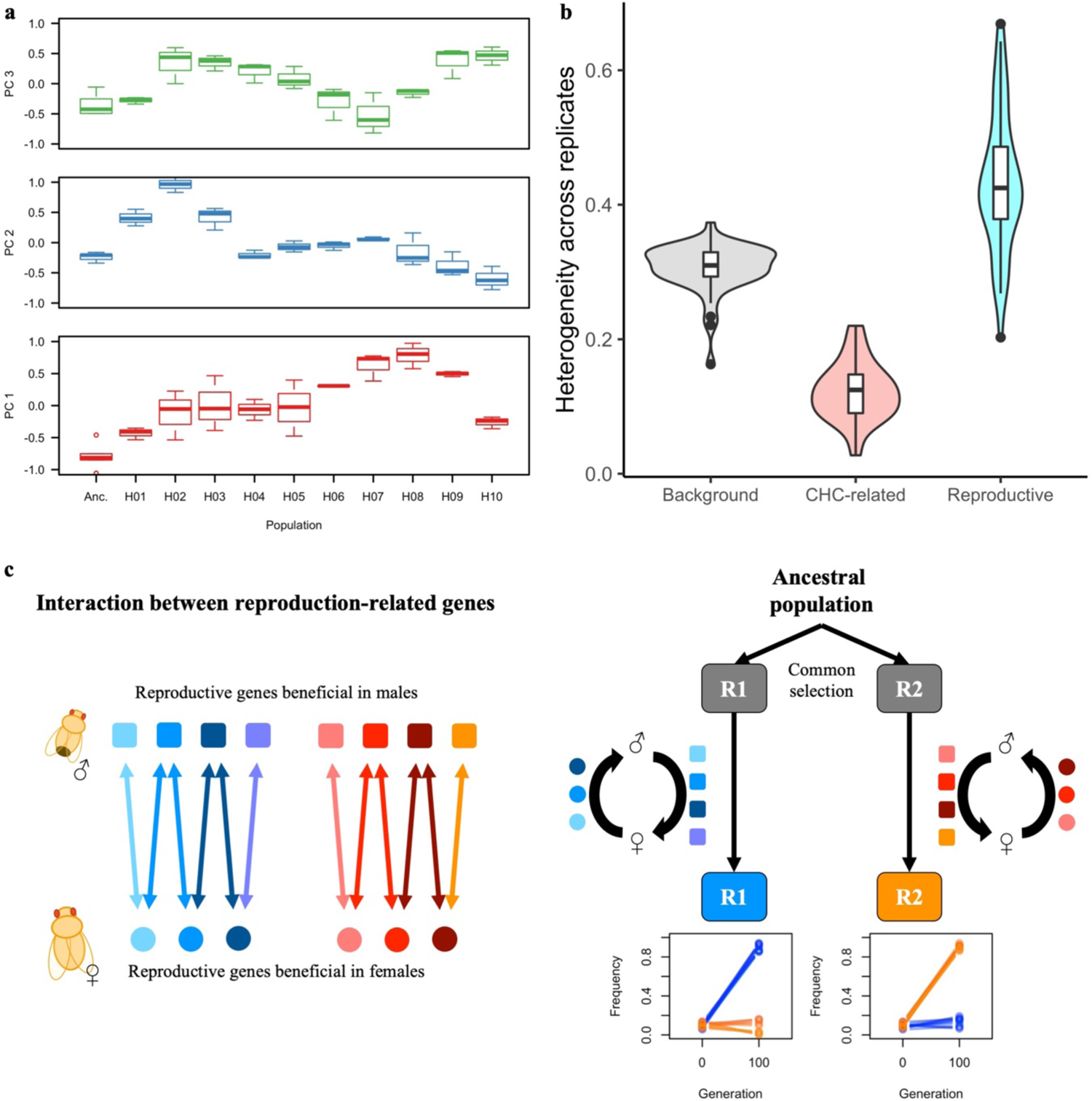
Heterogeneous adaptive changes in reproduction-related genes. **a.** Replicate-specific evolution caused the expression divergence of reproduction-related genes. The expression variance among 255 reproduction-related genes (see materials and methods) is summarized by the leading principal components (PCs). The first three PCs suggest that different replicates evolved different sets of genes during adaptation. **b.** The heterogeneity in expression evolution differs among different sets of genes: background (all expressed genes), CHC-related (33 genes reported in (Dembeck et al., 2015)), reproduction-related (255 genes from GO term (GO:0032504)). The heterogeneity is measured as *∑_i,j_(1 - cor(x_i_, x_j_))/N*, where x is a vector consisting of the expression of each gene in each evolution replicate; *i, j ∈ [1,10], i ≠ j* and N is the total number of pairwise combination of 10 replicates. Reproduction-related genes evolved with significantly higher heterogeneity than background genes or CHC-related genes (Kruskal-Wallis test, p < 0.001). **c.** An illustration for how intersexual co-evolution could lead to heterogeneous and incompatible evolution in isolated populations: Genes involved in male reproduction also serve important functions in females via either synergistic or antagonistic interaction with female reproduction-related genes. Multiple epistatic modules with different genes involved (red- and blue-colors) may co-exist in the ancestral population. A cascade of intersexual co-evolution for a certain group of genes is expected if selection pressure is imposed on either sex. As such co-evolutionary cascades (modules) are not unique, but redundant, replicate populations may evolve for high divergence and the alternative co-evolutionary paths may not be fully compatible with each other, resulting in fitness costs for between-replicate crosses.

## Discussion

The analysis of replicate populations adapting to the same environment demonstrated that reproductive isolation could rapidly evolve during adaptation to a novel temperature regime. The observed patterns of premating reproductive isolation were consistent with the predictions for ecological speciation - populations evolved to different habitats showed reproductive isolation, but no difference was detected between populations adapted to the same environment. Connecting the adaptive response in lipid metabolism of hot evolved flies to shifts in CHC profiles provides a plausible link between the adaptive response and the pleiotropic byproduct of reproductive isolation. Hence, our study presents one example of ecological adaptation where the target of selection (lipid metabolism) has been identified and the pleiotropic effects of involved genes explain the premating reproductive isolation.

Remarkably, we did not only find evidence for ecological speciation, but also for mutation-order speciation in the same experiment. While it has been previously proposed that inter-locus antagonistic coevolution could drive speciation [2, 27] and the decoupling of the coevolution of males and females has fitness consequences [30], the underlying molecular processes have remained unclear. Our study provides an important step forward. By showing that reproductive isolation between replicates is accompanied by an elevated variance in gene expression evolution of reproduction-associated genes, we provide a plausible explanation for the previously indirectly inferred coevolution between both sexes. Importantly, several lines of evidence suggest that the differences between replicates are not caused by genetic drift: 1) the variance in gene expression evolution of reproduction-associated genes among replicates is higher than that of randomly selected genes (under random genetic drift no enrichment for a GO category would be expected). 2) most of the reproduction-associated genes changed their expression in the same direction (mostly up-regulation) during adaptation 3) the higher expression of reproduction-related genes is consistent with the increased reproductive activity of the evolved populations (Figure 2a and [19]). This upregulation pattern was specific for reproduction-related genes and could not found for other divergently expressed genes (Figure 5 — Supplementary Figure 5). 4) the post-mating isolation among replicate evolved population is stronger for pairs with more diverged gene expression. With the limitation of only three data points, this further supports that alternative selection responses in replicate populations are the underlying genetic mechanism (Figure 5 — Supplementary Figure 6). It is not clear, however, if the higher reproductive activity of the hot-evolved flies is a direct adaptive response to temperature or reflects an adaptation to laboratory conditions.

Mutation-order speciation assumes that independent new mutations are fixed in replicate populations adapting to the same environment [6]. In our experiment, however, adaptation is probably driven by standing genetic variation [31]. Given that most traits are polygenic [32], it is conceivable that standing genetic variation is more important for speciation processes than previously thought. Polygenic control and heterogeneous genomic divergence have been documented in some iconic cases of replicated ecological speciation in nature such as the convergent morphological radiation of cichlid species in East African lakes [33, 34] as well as the repeated colonization to fresh water systems of oceanic stickleback species [35, 36]. We propose to expand the concept of mutation-order speciation by relaxing the condition of novel mutations. Potential incompatibilities may evolve among sister species with the same adaptive phenotypes, but alternative genetic pathways.

We observed the signature of mutation-order speciation from standing genetic variation in an experimental evolution study in the laboratory, but the important question is whether similar processes are likely to occur in natural populations. More work, both theoretical and empirical, is needed to determine the importance of mutation order speciation form standing genetic variation in a natural system.

## Material and Methods

### Experimental evolution

The setup and maintenance of the experimental populations are detailed in [37]. In brief, 10 replicated outbred populations were constituted from the same 202 isofemale lines derived from natural *Drosophila simulans* populations collected in Florida, USA in 2010. Five females of each line were pooled to create one replicate of a reconstituted ancestral population (henceforth called ancestral population). Replicated populations independently adapted to a laboratory environment at 18/28°C with 12hr dark/12hr light photoperiod with the census population size of 1250 adult individuals per population per generation Based on genomic allele frequency changes, an effective population size of ∼300 was inferred [37]. During the evolution experiment, all replicate populations were reared in the same incubator with randomization orientation/position across generations.

Each founder isofemale lines was maintained at small population size (typically less than 50 individuals) at 18 °C for >50 generations on standard laboratory food before the reconstitution of the ancestral populations for common garden experiments. Potential adaptation to the lab environment with the residual heterogeneity or de novo mutation is unlikely, as discussed previously [37]. The small effective population size during the maintenance of each line prevents adaptation to the lab environment. This has also been experimentally tested by the lack of significant difference in allele frequencies in populations which were reconstituted shortly after the establishment of the lines and after 50 generations [38]. Furthermore, mutations occurring in the laboratory, if any, are mostly recessive [39], and the effects are likely to be masked because after two generations of random mating during common garden maintenance, most individuals will be heterozygous for isofemale line-specific variants.

### Common garden experiment

We performed multiple common garden experiments with the evolved populations and reconstituted ancestral populations for the transcriptomic, metabolomic and phenotypic assays in this study. The common garden procedures have been detailed in [19, 20]. Briefly, replicates of ancestral populations were reconstituted by pooling five females each from the founder isofemale lines. These reconstituted ancestral populations and evolved replicates were reared at 18/28°C cycling with 12hr dark/12hr light photoperiod for at least two generations to minimize transgenerational and/or environmental effects before subjected to different phenotypic assays.

### Assay for male reproductive activity

We measured the reproductive activity for evolved and reconstituted ancestral populations at generation 140. After four generations at common garden condition (18/28°C cycling), 10 pairs of 5-day-old mated males and females were placed together in an agar-based arena (4% agar, 4% sugar) and filmed for 15 minutes at 20 FPS (frame-per-second) at 28°C using FlyCapture2 system (PointGrey, Version 2.13.3.31). In total, for matched-setup (flies from the same population are assayed together), eight trials within the evolved populations and six trials within the ancestral populations were assayed. In addition, five and six trials of reciprocal mismatched-setup (ancestral males were assayed together with evolved females and vice versa) were included in the experiment. Each fly was tracked using flytracker [40] for movement and behavior analysis. Janelia Automatic Animal Behavior Annotator (JAABA) was used to annotate and recognize the chasing behaviors [41]. The time a male fly spent on chasing females was quantified as its reproductive activity and the difference in the different setups was tested with a Kruskal-Wallis test in R.

### Multiple-choice mating assay for assortative mating

We performed a multiple-choice mating assay for assortative mating between 1) three ancestral populations and three selected evolved populations (repl. 1, 3 and 6, generation 194) 2) across the three independently evolved populations. We modified the experimental protocol from [42]. After two generations of common garden condition (18/28°C cycling with 12hr dark/12hr light photoperiod), 400 unmated males and females were collected from each population in six hours after eclosure. The collected flies were reared on corn meal in vials with moderate density (50 flies per vial, 8 replicates per sex per population) for four days and transferred onto yeast paste saturated with food coloring dyes (red or blue) one day before assaying. During the mating assay, 50 flies of each sex from each population in a combination of interest (one ancestral and one evolved populations/two evolved populations) were placed together in a cylindrical cage with a sealable entrance for aspirator. Eight cages (four for each color) for each population combination were setup and run simultaneously and each cage was checked sequentially. Mating pairs were captured with aspirator and identified under microscope based on the color of abdomen. The examination was terminated when the first 25 mating pairs in a cage were identified or when two hours elapsed. Based on a two-by-two contingency table from the assay for each population combination, Yule’s index (Y) [43] was calculated to quantify the strength of assortative mating and Fisher’s exact test was applied for hypothesis testing.

### Quantification and identification of cuticle hydrocarbons

We measured the cuticle hydrocarbons (CHCs) for five replicates of the ancestral population and 10 independently evolved populations at generation 158. After two generations in a common garden, three replicates of 20 virgin flies of each sex were collected two hours after eclosure from each population. The collected flies were aged for three additional days before CHC extraction. We used 100 !l of heptane containing an internal standard (IS, 10 !g n-C30) to wash the CHCs from each sample. The extracts were stored at −20°C until gas chromatography. GC/MS analysis was performed to isolate and quantify each compound. The analysis of chromatograph was performed with an Agilent 7890C gas chromatography system coupled to an Agilent 5975C MSD. Injection volume was 1 µl in an inlet at a temperature of 320°C, a split of 10:1. He was used as carrier gas at a velocity of 1 ml/min and the compounds separated in a HP-5MS column (30 m x 250 µm ID x 0.25 µm film thickness) with a temperature program 180°C to 240°C with 6°C/min. and then with 20°C/min. to 320°C. Transfer line temperature to the MSD was 280 °C, mass range of the MSD was 40-400 m/z and electromagnetic voltage was set at 70 eV.

Identification of the constituents were done based on retention index (RI), determined with reference to homologues series of n-alkanes (C8-C30) and mass spectra with the databases NIST 05. Each identified chemical compounds were validated with published *Drosophila* CHCs chromatograph (Dembeck et al., 2015; Pardy et al., 2019). For integration, identification and quantification of compounds, Automated Mass spectral Deconvolution and Identification System (AMDIS) [44] and openChrom [45] were used. The relative concentration of each CHC compound (excluding cis-Vaccenyl acetate) was calculated and subjected to centered-log ratio transformation [44]. Two-way analysis of variance (ANOVA) and principal component analysis (PCA) were performed to dissect the effect of evolution and sex on different CHCs.

### Cross-replicate compatibility assay

At generation 194 of the evolved populations, we performed a diallel cross among three selected evolved replicates (repl. 1, 3 and 6) and measured the number of progenies that survived until adulthood to approximate the fitness of each cross. Since we did not find assortative mating (Figure 2a), this assay can identify fitness differences caused by viability or fertilization success. The ancestral populations were not included because of the potential impact of assortative mating. After three generations of common garden rearing (18/28°C cycling), 75 unmated males and females were collected from each evolved replicate population and aged to four days old. For each cross, five males and five females from the same/different population were placed together in a vial (with food surface area of 3.14cm^2^; no crowding effect is expected) at the same common garden condition and two transfers were made for all vials with a 48-hour interval. The measurements of viable progeny numbers after 14 days from both transfers were averaged and treated as one replicate for the data analysis. Five replicates were generated for each cross and there were in total nine types of cross in a full diallel cross design among three populations.

### RNA-Seq data analysis for divergently expressed genes

RNA-Seq data for the evolution experiment at generation 103 were obtained from previous studies where contrasts between ancestral and evolved populations were made [19, 20]. In this study, we reanalyzed the data for the 5-day-old whole-body male samples of the evolved populations to identify the genes that were divergent across replicates. Only two replicates are available from each evolution replicates for females, preventing similar analysis in females [19, 46]. Genes with count per million (CPM) > 0.1 across all samples were retained and we modeled their expression as: *Y = repl + ε*, where *Y* is the normalized expression values; *repl* indicates the effect among evolution replicates and) is the random error. Likelihood ratio tests implemented in edgeR [47] were used to perform differential expression analysis on the effect of *repl*. Benjamini and Hochberg’s FDR correction [48] was applied with a FDR cutoff of 0.05. Genes differentially expressed in at least one replicate were identified for further analysis.

### Gene set enrichment analysis

In order to explore the functional implication of the parallel changes and random divergence in transcriptome, we tested for an enrichment of gene ontology (GO), tissue-specific expression and reproductive functions among the genes evolving for a parallel response [19] or those evolved for significant divergence across the 10 evolution replicates (this study). GO enrichment was performed using default “weight01” algorithm implemented in topGO (version 2.32.0) [49]. Genes highly expressed in each tissue were identified based on flyatlas2 expression dataset [50] (required > 2 fold higher expression in a certain tissue than whole-body). Genes involved in the cuticle hydrocarbon (CHC) metabolism were obtained from [25]. Fisher’s exact tests were applied for the enrichment analysis. Except for the GO enrichment analysis which already accounts for multiple testing [51], Benjamini and Hochberg’s FDR correction [48] was applied to account for multiple testing.

### Replicate-specific evolution in divergently evolving reproductive genes

In order to understand the driving force underlying the divergent evolution in the expression of genes involved in “multicellular organism reproduction” (GO:0032504), we investigated the replicate-specific gene expression evolution of 255 reproductive genes in this term that were significantly diverged across independent replicates. Using the expression matrix of these genes in all samples (Figure 5 — Figure Supplement 2), we summarized the variation in expression of the 255 genes by the principal components (PCs). The PC scores of each sample reflect the expression difference across the sample. Thus, comparing different evolved replicates to the ancestral samples could inform us about replicate-specific evolution in a certain subset of genes. Additionally, we investigated the heterogeneity in the expression changes across replicates as:

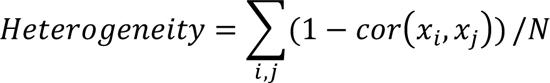

Where x is a vector consist of the expression change of each gene in each evolution replicate, *i,j ∈ [1,10], i ≠ j* and N is the total number of pairwise combination of 10 replicates.

### Data and materials availability

Sequence reads from this study is available from the European Sequence Read Archive (http://www.ebi.ac.uk/ena/) under the study accession number PRJEB35504 and PRJEB35506. Additional scripts and raw data are available on Github upon publication.

